# The evolution of the age of onset of resistance to infectious disease

**DOI:** 10.1101/2022.11.01.514666

**Authors:** Lydia J. Buckingham, Ben Ashby

## Abstract

Many organisms experience an increase in disease resistance as they age but the time of life at which this change occurs varies. Increases in resistance are partially due to prior exposure and physiological constraints but these cannot fully explain the observed patterns of age-related resistance. An alternative explanation is that developing resistance at an earlier age incurs costs to other life-history traits. Here, we explore how trade-offs with host reproduction or mortality affect the evolution of the onset of resistance, depending on when during the host’s life-cycle the costs are paid (only when resistance is developing, only when resistant or throughout the lifetime). We find that the timing of the costs is crucial to determining evolutionary outcomes, often making the difference between resistance developing at an early or late age. Accurate modelling of biological systems therefore relies on knowing not only the shape of trade-offs but also when they take effect. We also find that the evolution of the rate of onset of resistance can result in evolutionary branching. This provides an alternative, possible evolutionary history of populations which are dimorphic in disease resistance, where the rate of onset of resistance has diversified rather than the level of resistance.

## Introduction

Many organisms experience changes in their level of resistance to infectious disease as they age, with repercussions for both their own health and transmission to others (Altizer et al. 2004; Apolloni et al. 2013; Clark et al. 2017). Age-related differences in resistance have been recorded across many taxa, including plants (Miller 1983; Panter and Jones 2002; Develey-Rivière and Galiana 2007; Bruns et al. 2017), invertebrates (Sait et al. 1994; Kubi et al. 2006; Armitage and Boomsma 2010; Garbutt et al. 2014; Green et al. 2016; Klemme et al. 2022) and vertebrates (Duca 1948; Zuckerman and Yoeli 1954; Francis 1961), including humans (Baird 1998; Kurtis et al. 2001; Glynn and Moss 2020). In particular, adults are generally more resistant than juveniles (Duca 1948; Zuckerman and Yoeli 1954; Francis 1961; Sait et al. 1994; Baird 1998; Kurtis et al. 2001; Panter and Jones 2002; Kubi et al. 2006; Develey-Rivière and Galiana 2007; Armitage and Boomsma 2010; Green et al. 2016; Bruns et al. 2017), with some species being completely resistant as adults to pathogens which are highly detrimental to juveniles (e.g. nucleopolyhedrovirus in gypsy moths (Elderd et al. 2008)). Precisely when in the host’s lifetime these changes in resistance occur varies between systems. This variation in the timing of resistance onset has received little empirical or theoretical attention, with very few studies seeking to explain or even describe the time within a host’s lifespan at which resistance changes. A key outstanding question is therefore, ‘how should evolution shape the timing of the onset of resistance?’

As there is variation in the age of onset of resistance within species and resistance traits are often heritable, it is reasonable to assume that the age of onset of resistance is an evolvable trait. In this paper, we seek to generate predictions for how the age of onset of resistance might evolve. Intuitively, an earlier onset of resistance will tend to lower disease prevalence and hence reduce the risk of infection, thereby acting as a negative feedback on selection if resistance is costly. However, precisely how this phenomenon affects the evolution of the onset of resistance remains to be determined.

Several factors are known to contribute towards the onset of disease resistance. Many species’ immune systems respond to prior exposure, meaning that resistance may increase substantially following infection. This mechanism can explain some, but not all, observed variation in the age of onset of disease resistance, particularly in the case of species which rely solely or predominantly on innate resistance. For instance, different species of plants experience changes in resistance at different life-stages (Develey-Rivière and Galiana 2007). Alternatively, variation in the age of onset of resistance may have evolved due to contrasting selection pressures at different life-stages. For instance, if a pathogen is more harmful to older hosts than younger hosts then the onset of resistance may be delayed if developing or maintaining resistance is costly. In some cases, however, age-related changes in resistance have been observed towards pathogens which have similar effects on hosts of all ages (Bruns et al. 2017), suggesting that the timing of the onset of resistance is also affected by other factors. Hence, none of these mechanisms can provide a full explanation for variation in the age of onset of disease resistance.

An alternative evolutionary explanation for increases in resistance with age is that heightened resistance incurs costs to host life-history traits. For instance, hosts which invest in developing resistance quickly may allocate resources away from growth or development which may in turn impact on future reproduction or survival. In principle, these reductions in fecundity or survival may only be experienced while resistance is developing but such costs may also have long-term consequences which affect the reproduction or survival of the host later in life (Chaplin and Mann 1978; Simons 1979; Tian et al. 2003). Evolutionary trade-offs may therefore provide an explanation as to why hosts might develop resistance later in life rather than being resistant from birth or from a very early age. Empirical studies reveal trade-offs between reproduction (Biere and Antonovics 1996; Karasov et al. 2017) or growth (Biere and Antonovics 1996; Susi and Laine 2015; Karasov et al. 2017; Bartlett et al. 2018) and disease resistance in plants and insects, but little data is available on age-specific costs (Izhar and Ben-Ami 2015).

Epidemiological models have considered disease spread in age-structured populations (e.g. (Clark et al. 2017)) and evolutionary models have considered the evolution of resistance in populations with no age structure (e.g. (Antonovics and Thrall 1994; Boots and Haraguchi 1999; Carlsson-Graner and Thrall 2006; Miller et al. 2007; Donnelly et al. 2015)). In contrast, the evolution of age-structured resistance has received relatively little attention. However, recent theory has explored the evolution of age-specific resistance in juveniles and adults, showing that differences in resistance between life-stages are most pronounced for particularly long- or short-lived hosts (Ashby and Bruns 2018) and that adult resistance often evolves to exceed juvenile resistance, depending on which traits trade off with resistance (Buckingham et al. 2023). Here, we investigate a different characteristic of age-specific resistance, namely the timing of the onset of resistance. We focus our analysis on trade-offs with mortality and reproduction, which may affect the host during development of resistance, during the resistant stage or throughout the lifetime of the host. We also consider variation in pathogen traits, specifically transmissibility and the strength and type of virulence. We find that the timing of the trade-offs (costs paid before or after the onset of resistance or throughout the host’s lifetime) has a significant effect on evolutionary outcomes, often determining whether the host becomes resistant very early or very late in life. We also find that evolutionary branching of the age of onset of resistance can generate a dimorphic population where some hosts are resistant throughout their lives and others never develop resistance.

## Methods

### Model description

We consider a well-mixed, asexual host population in which hosts are born fully susceptible to an infectious disease but develop full innate resistance to infection at a constant (but evolvable) rate *ζ* > 0 (see Fig. 1A for a model schematic and Table 1 for a full list of parameters and variables). The length of time before an individual becomes resistant is therefore exponentially distributed (with parameter 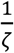) and so 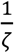 gives the average age of onset of resistance. Let *S, I* and *R* be the densities of susceptible, infected and resistant hosts respectively, giving a total host population density of *N* = *S* + *I* + *R*. Non-resistant hosts reproduce at a maximum rate *a* > 0, subject to density-dependent competition given by *q* > 0, and die naturally at rate *b* > 0. Similarly, resistant hosts reproduce at rate *a_R_* and die naturally at rate *b_R_*. Pathogen transmission is assumed to be density-dependent, with transmission rate *β*, and infected hosts may either experience sterility virulence equal to 1 – *f*, where 0 ≤ *f* ≤ 1 is the reduction in fecundity when infected, or mortality virulence given by *α* > 0, the relative increase in the mortality rate. We assume that hosts do not recover from infection, as recovery would complicate the timing of the onset of resistance (it is not clear how infected individuals would transition to the innately resistant stage in this case) and would also require us to account for potential long-term effects of sterility virulence.

**Fig. 1.**
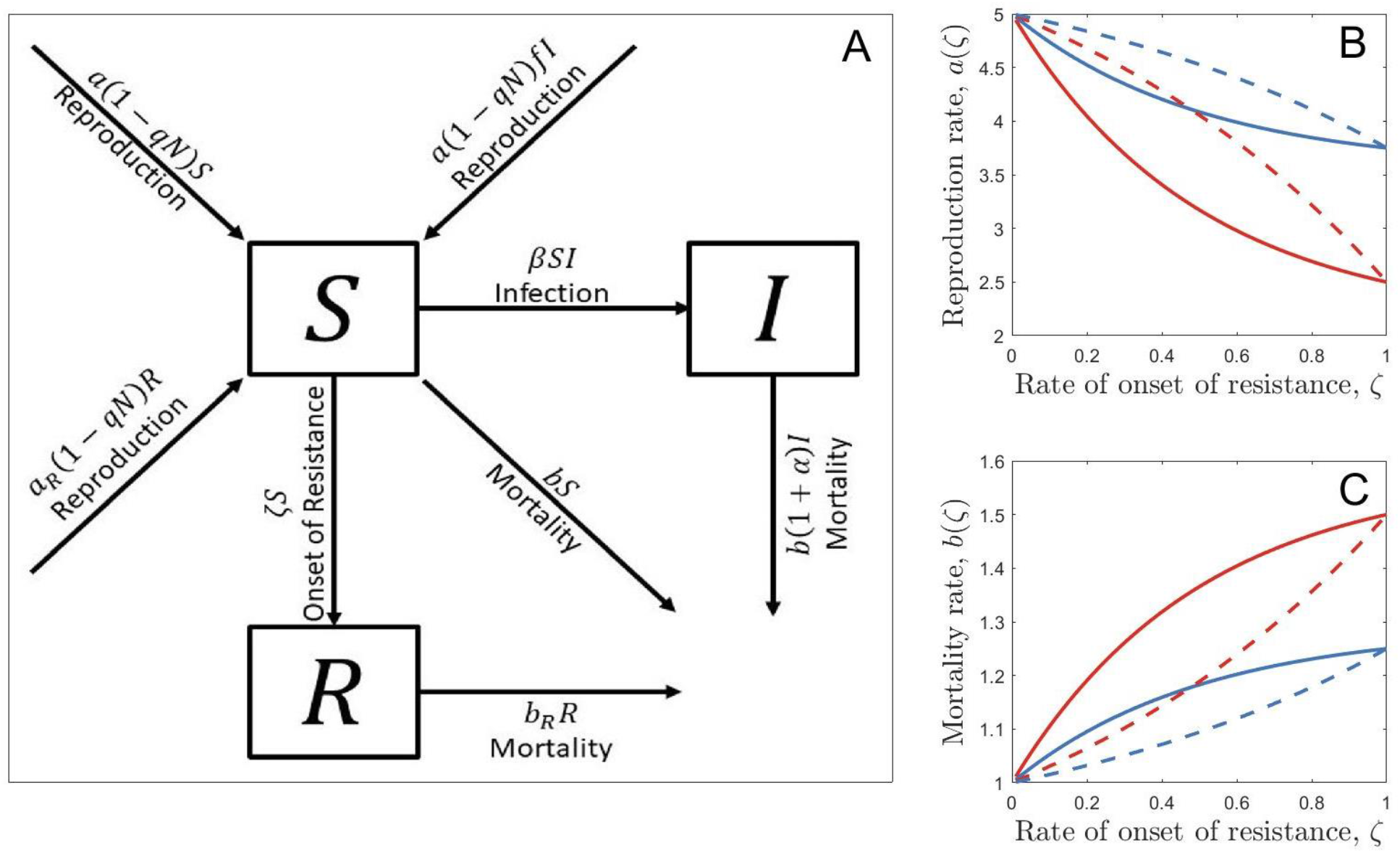
(A) Model schematic for a monomorphic population. (B) Examples of reproduction trade-off functions (with *a*_0_ = 5). (C) Examples of mortality trade-off functions (with *b*_0_ = 1). Trade-off strength is controlled by the parameter 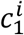; a relatively strong trade-off (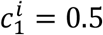, red) results in a much larger reduction in the birth rate for a given level of adult resistance than a relatively weak trade-off does (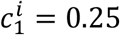, blue). Trade-off shape is controlled by the parameter 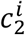; a positive value (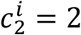, solid) means that the costs of resistance decelerate (increasing returns) as the rate of onset gets faster whereas a negative value (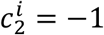, dashed) leads to accelerating costs (diminishing returns).

**Table 1.** Descriptions and default values or ranges of model parameters and variables.

| Parameter/<br>variable | Description | Default value<br>or range |
| --- | --- | --- |
| $a, b$ | Reproduction rate/natural mortality rate of non-resistant hosts | n/a |
| $a_R, b_R$ | Reproduction rate/natural mortality rate of resistant hosts | n/a |
| $a_0$ | Baseline value of the reproduction rate | 5 |
| $b_0$ | Baseline value of the natural mortality rate | 0.1 |
| $c_1^a, c_1^b$ | Strength of reproduction/mortality trade-offs | 0.1 |
| $c_2^a, c_2^b$ | Shape of reproduction/mortality trade-offs | $\pm 2$ |
| $1 - f$ | Sterility virulence | $0 \leq f \leq 1$ |
| $\zeta$ | Rate of onset of resistance | $0 \leq \zeta$ |
| $S, I, R$ | Density of susceptible/infected/resistant hosts | n/a |
| $N$ | Total host population density | n/a |
| $q$ | Strength of host density dependence | 1 |
| $t$ | Time, measured in arbitrary units | n/a |
| $\alpha$ | Mortality virulence (proportional increase in mortality rate) | $0 \leq \alpha$ |
| $\beta$ | Pathogen transmission rate | 0.5 |
| $1 - h$ | Relative reduction in fecundity when resistant | $0 \leq h \leq 1$ |
| $\delta$ | Relative increase in mortality when resistant | $0 \leq \delta$ |

In a monomorphic population and in the absence of any costs of resistance, the population dynamics are described by the following set of ordinary differential equations:

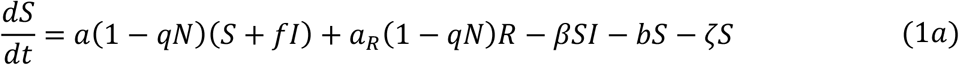

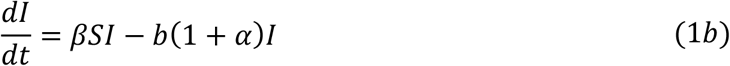

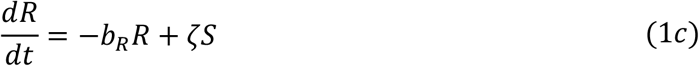

Intuitively, in the absence of any costs of resistance, selection will favour ever larger rate of onset of resistance (*ζ*) so that there is an immediate onset of resistance at the start of the host lifespan, which will in turn drive the pathogen extinct. We therefore consider cases where being resistant or developing resistance incurs a cost to the host, either through higher mortality or lower fecundity (e.g., due to resource allocation). Such costs might be paid only when the host is resistant (i.e., in the *R* class), or prior to the onset of resistance (i.e., during development in classes *S* and *I*), or throughout the lifetime of the host. The magnitude of any costs may therefore be constant, or depend on the rate of onset of resistance, *ζ*. To account for costs of resistance, we modify equation (1) so that the reproduction rates, *a*(*ζ*) and *a_R_*(*ζ*), and mortality rates, *b*(*ζ*) and *b_R_*(*ζ*), may depend on the rate of onset of resistance.

We consider six possible scenarios:

1. A constant fecundity cost, paid only when the host is resistant (*a_R_*(*ζ*) = *ah*, where 1 – *h* is the relative reduction in fecundity when resistant, with *b_R_*(*ζ*) = *b, a* = *a*_0_ and *b* = *b*_0_)
2. A constant mortality cost, paid only when the host is resistant (*b_R_*(*ζ*) = *b*(1 + *δ*), where *δ* is the relative increase in mortality when resistant, with *a_R_*(*ζ*) = *a*, *a* = *a*_0_ and *b* = *b*_0_)
3. A whole-life fecundity cost (*a_R_*(*ζ*) = *a*(*ζ*) as defined in equation (2a), with *b_R_*(*ζ*) = *b* and *b* = *b*_0_)
4. A whole-life mortality cost (*b_R_*(*ζ*) = *b*(*ζ*) as defined in equation (2b), with *a_R_*(*ζ*) = *a* and *a* = *a*_0_)
5. A developmental fecundity cost (*a*(*ζ*) as defined in equation (2a), with *a_R_*(*ζ*) = *a*_0_, *b_R_*(*ζ*) = *b* and *b* = *b*_0_)
6. A developmental mortality cost (*b*(*ζ*) as defined in equation (2b), with *a_R_*(*ζ*) = *a, b_R_*(*ζ*) = *b*_0_ and *a* = *a*_0_).

where *a*_0_ and *b*_0_ are baseline reproduction and mortality rates and we use the following trade-offs for the reproduction rate:

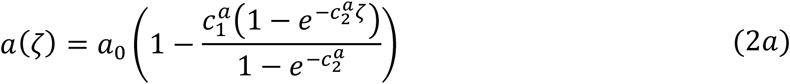

and the mortality rate:

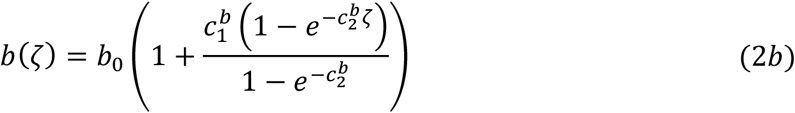

where 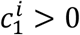 determines the maximum strength of the trade-off (i.e. the maximum proportional reduction or increase in the associated life-history trait) and 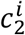 determines the shape of the trade-off (larger absolute values correspond to greater deviations from linearity; Fig. 1B & Fig. 1C). Note that it is possible to rescale the system of equations (1a) to (1c) so that we can set *q* = 1 and *b*_0_ = 1 without loss of generality (see *Supplementary Materials*).

### Epidemiological dynamics

The disease-free equilibrium of system (1) is given by:

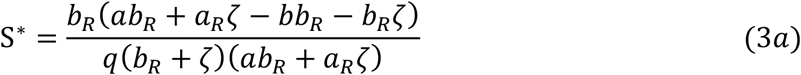

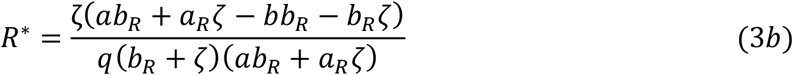

and is stable provided that *ab_R_* + *a_R_ζ* > *bb_R_* + *b_R_ζ* and

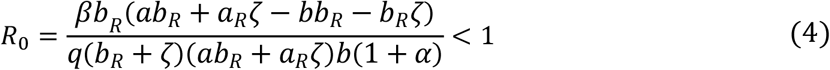

where *R*_0_ is the basic reproductive ratio of the pathogen (see *Supplementary Materials* for derivation). The pathogen can spread when *R*_0_ > 1, in which case there is a stable, endemic (non-trivial) equilibrium (see *Supplementary Materials*).

### Evolutionary invasion analysis

We use evolutionary invasion analysis (adaptive dynamics) to determine the evolutionary dynamics of the rate of onset of resistance, *ζ* (Geritz et al. 1998). Specifically, we assume that mutations are sufficiently rare that there is a separation of ecological and evolutionary timescales (the ecological dynamics of the resident population reach equilibrium before the next mutation occurs) and are sufficiently small that *ζ_m_* ≈ *ζ*, where *ζ_m_* is the mutant trait and *ζ* is the resident trait. The invasion dynamics of rare host mutants in a resident population at its endemic equilibrium are given by:

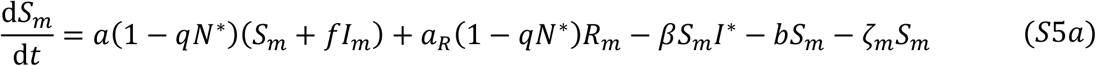

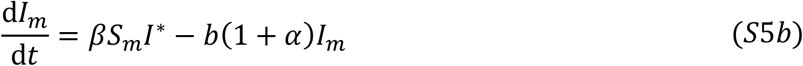

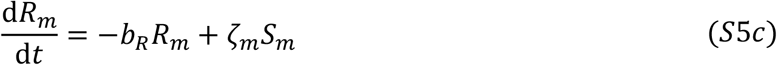

where asterisks denote the endemic equilibrium of the resident population and *a, a_R_, b* and *b_R_* may be functions of *ζ_m_*, depending on the nature of the costs of resistance.

Using the next generation method (Hurford et al. 2010), one can derive the following expression for the invasion fitness of a rare mutant in a resident population (see *Supplementary Materials*):

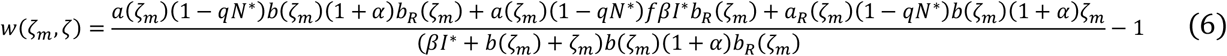

In the general case, the lengthy expression for the endemic equilibrium limits analytical progress. However, in cases (1) and (2), where the costs of resistance are constant, we can proceed to determine an expression for the singular value (value which maximises or minimises the invasion fitness) of the rate of onset of resistance.

### Simulations

We can also use simulations to investigate the evolution of *ζ* (the rate of onset of resistance). To simulate an evolutionary trajectory of the evolving trait (*ζ*), we first choose a resident value for *ζ* and an initial composition for the host population. We then run the ecological dynamics for a fixed length of time, *T*. At this stage, we introduce a small, second sub-population with a different value of *ζ* (chosen at random to be either slightly higher or lower than the resident value) to model the introduction of a rare mutant into the population. The ecological dynamics (given for an arbitrary number of phenotypes in the *Supplementary Materials*) are then run again for this new population for a fixed length of time, *T*. If the size of the sub-population with a particular value of *ζ* falls below a threshold, then that sub-population is removed (it has gone extinct). An additional, small sub-population with a new value of *ζ* (chosen at random to be slightly higher or lower than one of the existing trait values in the population) is introduced at each evolutionary timestep, the ecological dynamics are run and sub-populations that fall below the extinction threshold are removed. This process is repeated for many evolutionary timesteps to simulate the evolution of *ζ*.

## Results

### Constant costs

We begin by considering the effects of constant fecundity and mortality costs (cases 1 and 2 described in the methods) on the evolution of the onset of resistance, *ζ*. This means that we have a constant fecundity cost when resistant *a_R_* = *ah*, where the reproduction rate when susceptible is *a* = *a*_0_, and a constant mortality cost when resistant, *b_R_* = *b*(1 + *δ*), where the mortality rate when susceptible is *b* = *b*_0_. In this case, we can differentiate the invasion fitness with respect to the mutant trait *ζ_m_* to calculate the fitness gradient (see *Supplementary Materials*) and then find the root of the fitness gradient to give us the singular strategy for the rate of onset of resistance:

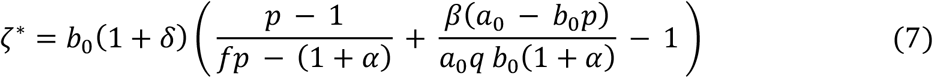

where 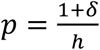 for notational convenience. Note that this expression can take a negative value, in which case the singular strategy is effectively at *ζ\** = 0 (no onset of resistance).

We can see from this expression that the rate of onset of resistance rises linearly as the pathogen transmissibility increases. This is because a more transmissible parasite will have a higher disease prevalence and so will exert stronger selection for fast onset of resistance. The singular strategy also rises as sterility virulence increases because more virulent pathogens impose stronger selection for resistance (Fig. 2D). This pattern is also seen as mortality virulence rises but only as long as mortality virulence is not too high. When mortality virulence is high, infected hosts die quickly, reducing the infectious period and so the potential for disease transmission. This reduces the pathogen density and so weakens selection for fast onset of resistance, leading to a fall in the rate of onset of resistance (Fig. 2B).

**Fig. 2.**
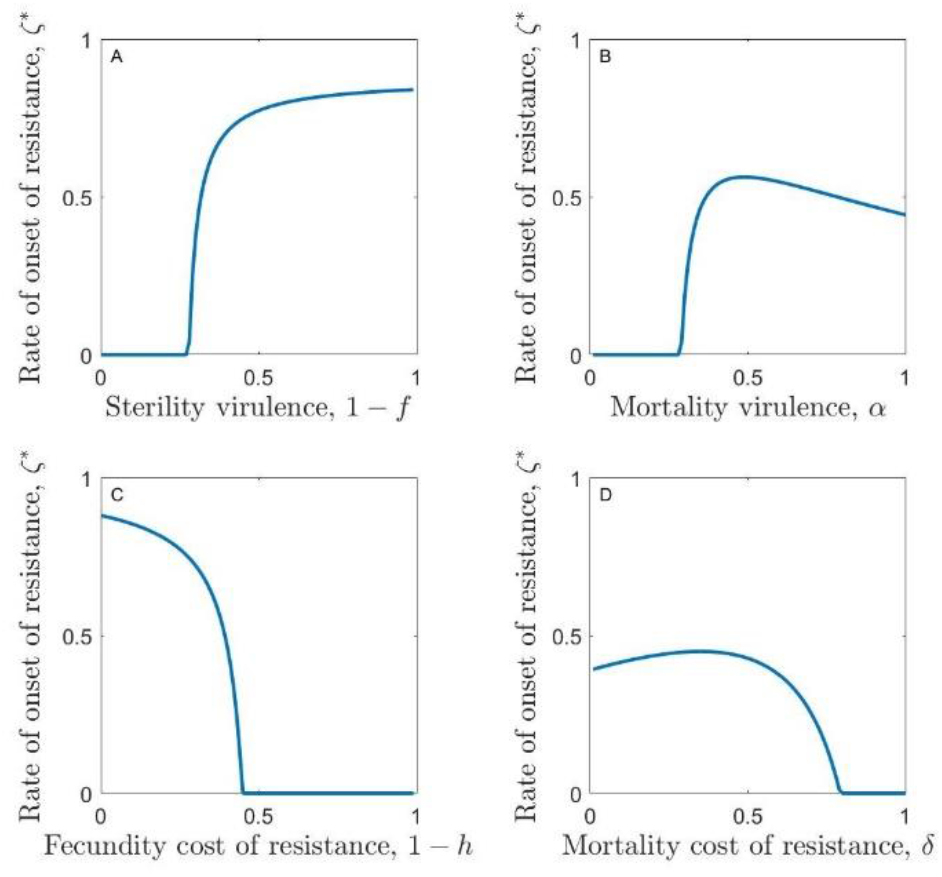
Evolution of the rate of onset of resistance, *ζ\**, in the case of constant costs of resistance. Parameters used are as in Table 1 except for *β* = 1, with (A) 1 – *f* = 0, 1 – *h* = 0 and *a* = 1, (B) 1 – *f* = 0, *1 – h = 0* and *δ* = 0.25, (C) 1 – *f* = 0.5, *a* = 0 and 5 = 0 and (D) 1 – *h* = 0.25, *a* = 0 and *δ* = 0.

We might also expect that the rate of onset of resistance would fall as the costs of being resistant rise. This is indeed true when the costs of fast onset of resistance are fecundity costs or when they are relatively large mortality costs (Fig. 2A, C). However, when low mortality costs are paid, it is possible for the rate of onset to rise as the costs of being resistant increase (Fig. 2A). This is because raising the mortality costs causes a significant increase in the density of the pathogen; resistant individuals die faster and so there are fewer of them in the population, yet reproduction is still high enough to maintain the population near to its density-dependent limit, meaning that the density of non-resistant individuals rises. This leads to a population with a reduced proportion of resistant individuals and an increased proportion of infected individuals. This heightened pathogen density then imposes greater selection for fast onset of resistance.

Analytical calculations reveal the invasion fitness of any mutant in a resident population at the singular strategy is zero: *w*(*ζ_m_, ζ*\*) = 0 (see *Supplementary Materials*). Hence 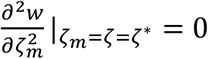 always and so the singular strategy is not strictly evolutionarily stable. The expression which determines convergence stability 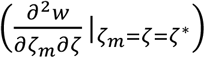 is intractable and so it is not possible to determine conditions for convergence stability analytically. However, extensive numerical calculations suggest that the singular strategy is always convergence stable (see *Supplementary Materials*). In order to classify the stability of the singular strategy fully however, we also need to determine whether it appears to be evolutionarily stable or unstable in practice. To do this, we use simulations. Evolutionary simulations (see *Supplementary Materials* for description) reveal that the singular strategies behave like continuously stable strategies (CSS’s), for a wide range of parameter sets. Fig. 3 depicts graphically the simulated changes in the rate of onset of resistance over evolutionary time, showing how it tends towards the analytically determined singular strategy and then remains there.

**Fig. 3.**
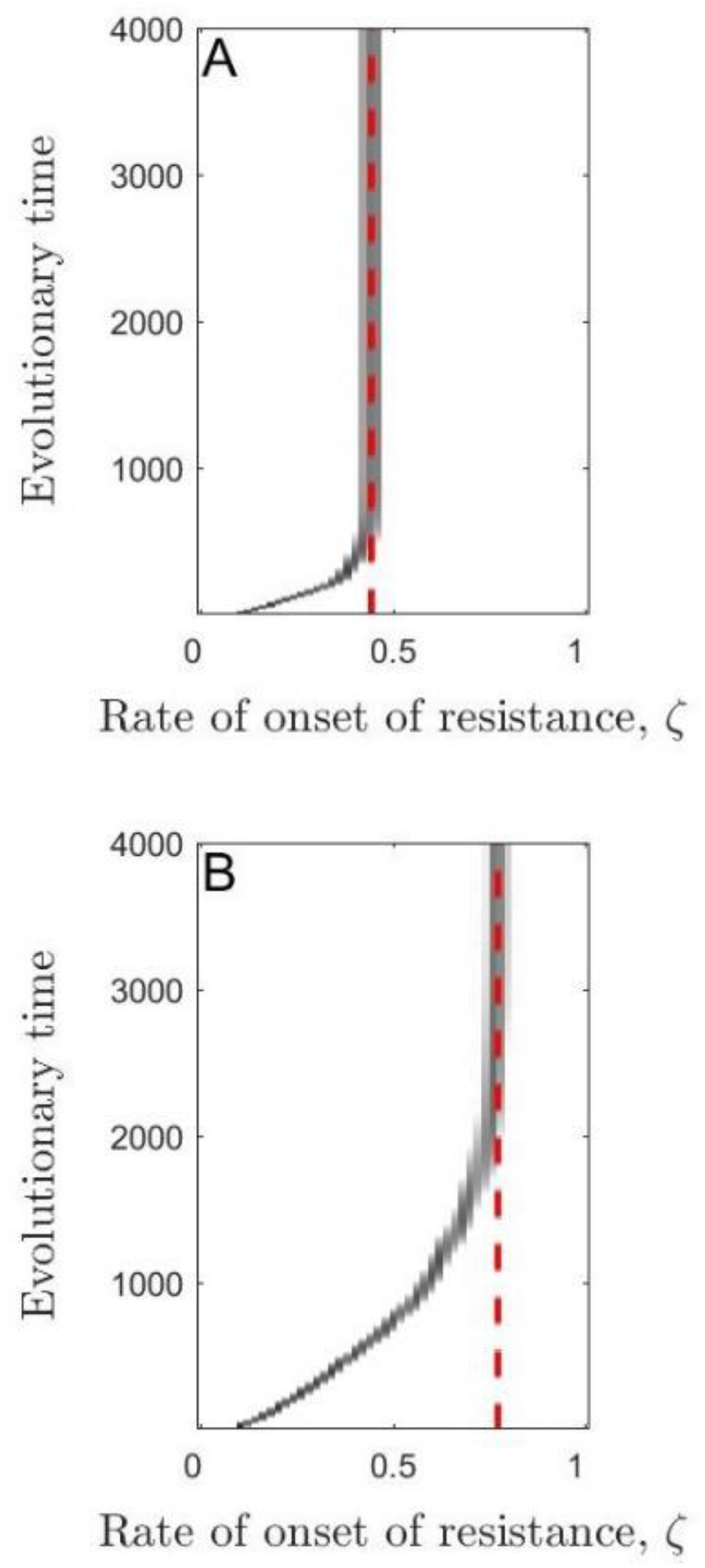
Simulated trajectories of the evolution of the rate of onset of resistance over time. The red, dashed lines show the analytically determined values of the singular strategies. Parameters used are as in Table 1 except for *β* = 1, with (A) 1 – *f* = 0, *α* = 1, 1 – *h* = 0 and *δ* = 0.25 in the case of scenario (2) where the host pays a constant mortality cost when resistant and with (B) 1 – *f* = 0.5, *α* = 0, 1 – *h* = 0.25 and *δ* = 0 in the case of scenario (1) where the host pays a constant fecundity cost when resistant.

### Non-constant costs

We now consider non-constant costs for the onset of resistance (cases 3-6 described in the methods). When costs to reproduction or mortality accelerate 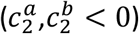, the rate of onset of resistance always evolves to a single, continuously stable strategy (CSS). When the costs decelerate, there are more possible evolutionary outcomes; a single CSS (the trait evolves towards this value and then remains there), repeller (the trait evolves away from this value, tending towards zero, another singular strategy, or increasing indefinitely) or branching point (the trait evolves towards this value but then undergoes disruptive selection and splits into two sub-populations) may be observed, as well as a repeller above a branching point, a branching point above a repeller or two repellers with a branching point in between (parameter values for which these occur are shown in Fig. S1 and Fig. S2).

When the costs of having a faster onset of resistance are paid throughout the host’s lifetime (cases 3 & 4), we can show analytically that the singular strategy (or the uppermost singular strategy if there is more than one) is always convergence stable (see *Supplementary Materials*). When the costs of having a faster onset of resistance are only paid before the onset of resistance (cases 5 & 6), this singular strategy is never convergence stable (see *Supplementary Materials*). This means that, if the rate of onset of resistance is initially sufficiently high, then it will increase indefinitely, in effect causing new-born hosts to become resistant immediately. This is because the host is already paying a significant cost for a reasonably fast onset of resistance. It is more beneficial for hosts to pay a slightly higher cost for a shorter time (before it becomes resistant and stops paying the cost) than to pay a slightly lower cost for a longer time.

Many factors influence whether the onset of resistance is fast or slow. However, certain parameters have especially strong effects. We can see these effects most clearly if we consider accelerating costs 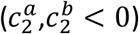, as each set of parameters produces a single evolutionary endpoint which does not depend on the initial conditions. We find that the virulence of the pathogen plays a particularly important role, with the fastest onset of resistance evolving when the pathogen exerts weak mortality virulence and strong sterility virulence (see Fig. 4; note that contours show the proportion of the host population which is resistant to the pathogen). This is because strong mortality virulence reduces disease prevalence by reducing the average duration of infection and so weakens selection for resistance. Sterility virulence, in contrast, does not affect the average duration of infection and so does not reduce disease prevalence but does greatly reduce host fitness. The fastest onset of resistance therefore evolves when disease prevalence is high (low mortality virulence) and when there is strong sterility virulence. Conversely, the rate of onset of resistance tends to zero when both sterility and mortality virulence are negligible (because the cost of infection is minimal) or when mortality virulence is very high (because disease prevalence is so low that the host is unlikely to be exposed to the pathogen). The results described above are qualitatively similar for decelerating trade-offs when the costs of resistance are paid throughout the lifetime of the hosts and there is a single CSS.

**Fig. 4.**
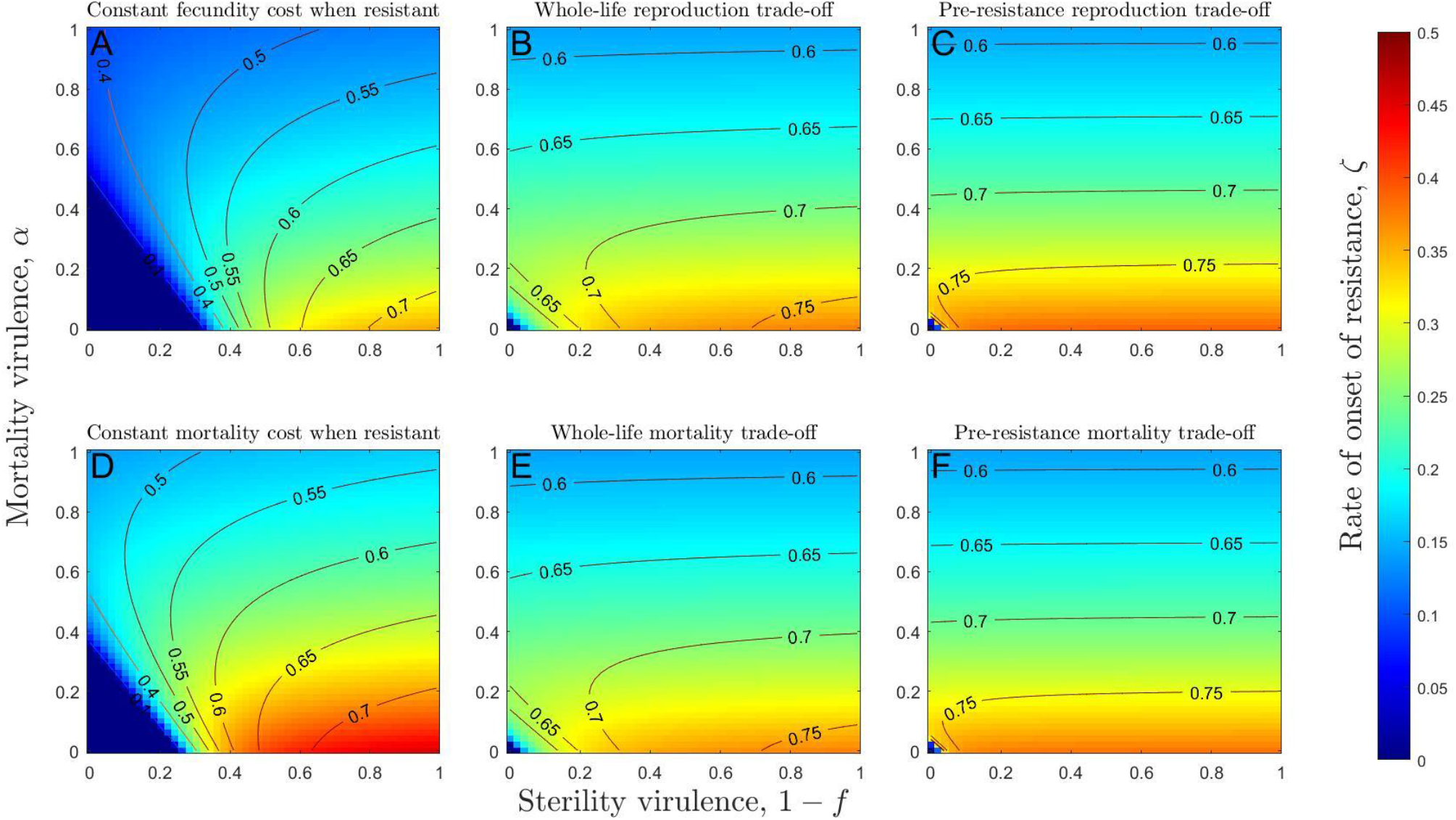
Heatmaps showing the evolved rate of onset of resistance as mortality and sterility virulence vary and for qualitatively different trade-offs. Contours show the proportion of the host population that is resistant to the disease and so do not correspond to heatmap colours. The rate of onset of resistance is highest (red) when mortality virulence is low and sterility virulence is high (bottom right of each panel). Increasing sterility virulence (left to right within each panel) always causes the rate of onset of resistance to increase. Increasing mortality virulence (bottom to top within each panel) may also cause the rate of onset of resistance to rise and then fall for sufficiently low values of sterility virulence. These patterns are broadly consistent across all six trade-off scenarios (A-F). Parameters used are as in Table 1, except 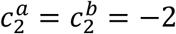 (accelerating costs), *h* = 0.75 and *δ* = 0.25, where applicable.

In general, greater pathogen transmissibility leads to the evolution of faster onset of resistance. However, as pathogen transmissibility gets very high, the rate of onset of resistance begins to fall (Fig. 5). This is because higher transmissibility corresponds to a lower average age on infection, and so it is more costly for a host to invest in a rapid onset of resistance (which will have to be very fast to take effect before the host becomes infected by a highly transmissible pathogen) than it is for the host to become infected with the pathogen (having not paid the costs of fast onset of resistance). This is consistent for both fecundity and mortality trade-offs (Fig. 5) of a variety of different strengths (values of 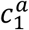 and 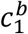). As expected, greater costs (higher values of 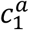 and 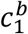) lead to the evolution of slower onset of resistance. Fecundity and mortality costs with the same proportional effects (e.g., a 50% increase in mortality or a 50% decrease in reproduction) have similar quantitative impacts on the rate of onset of resistance (Fig. 5).

**Fig. 5.**
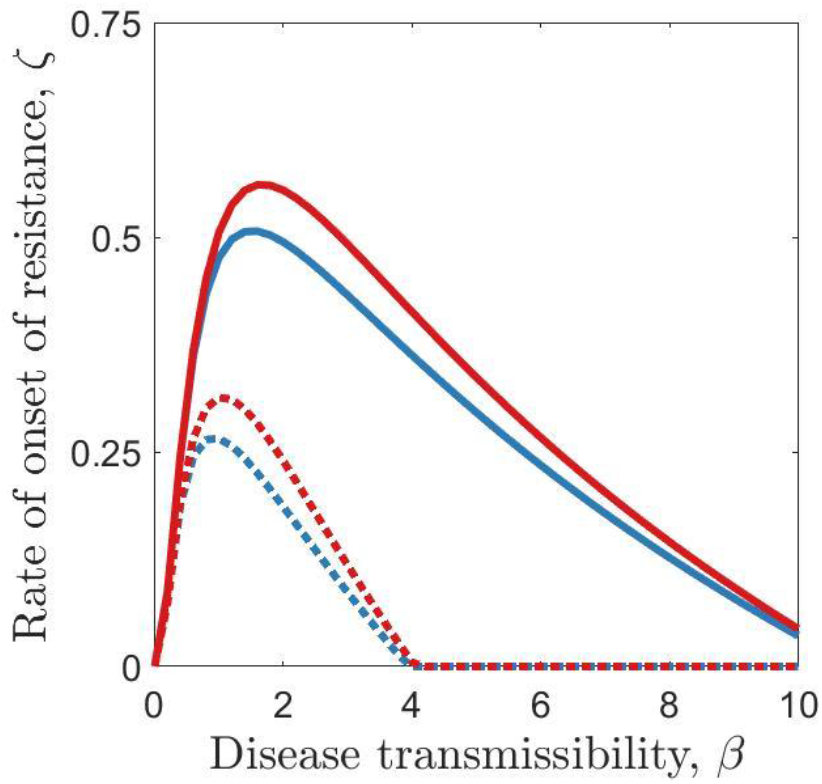
The effect of disease transmissibility (*β*) on the rate of onset of resistance, when either a fecundity cost (blue curves) or a mortality cost (red curves) is paid throughout the lifetime of the host (scenarios 3 and 4). The qualitative effects of transmissibility are the same whether the costs of fast onset of resistance are low (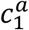 or 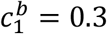; solid) or high (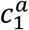 or 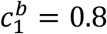; dotted). Parameters used are as in Table 1 with 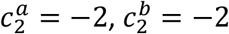, *f* = 0.5 and *a* = 0.

### Diversification

The host population may diversify through evolutionary branching when the costs are decelerating (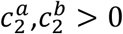; see Fig. 6). Evolutionary branching leads to one sub-population with a very high rate of onset of resistance (*ζ* large) that increases indefinitely (see *Supplementary Materials*), until these individuals effectively develop resistance immediately after birth, and another sub-population with a very low rate of onset of resistance (*ζ* ≈ 0), with these individuals unlikely ever to become resistant given the typical lifespan of the host. When costs are paid throughout the lifetime of the host, these costs must be strongly decelerating 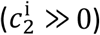 for branching to occur (Fig. 7A). When costs are paid before the onset of resistance, branching only occurs for weakly decelerating costs (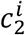 small but positive). The latter branching points always sit below repellers and so the population will only branch if the initial rate of onset of resistance is sufficiently low (otherwise, the rate of onset of resistance will increase indefinitely; Fig. 7B).

**Fig. 6.**
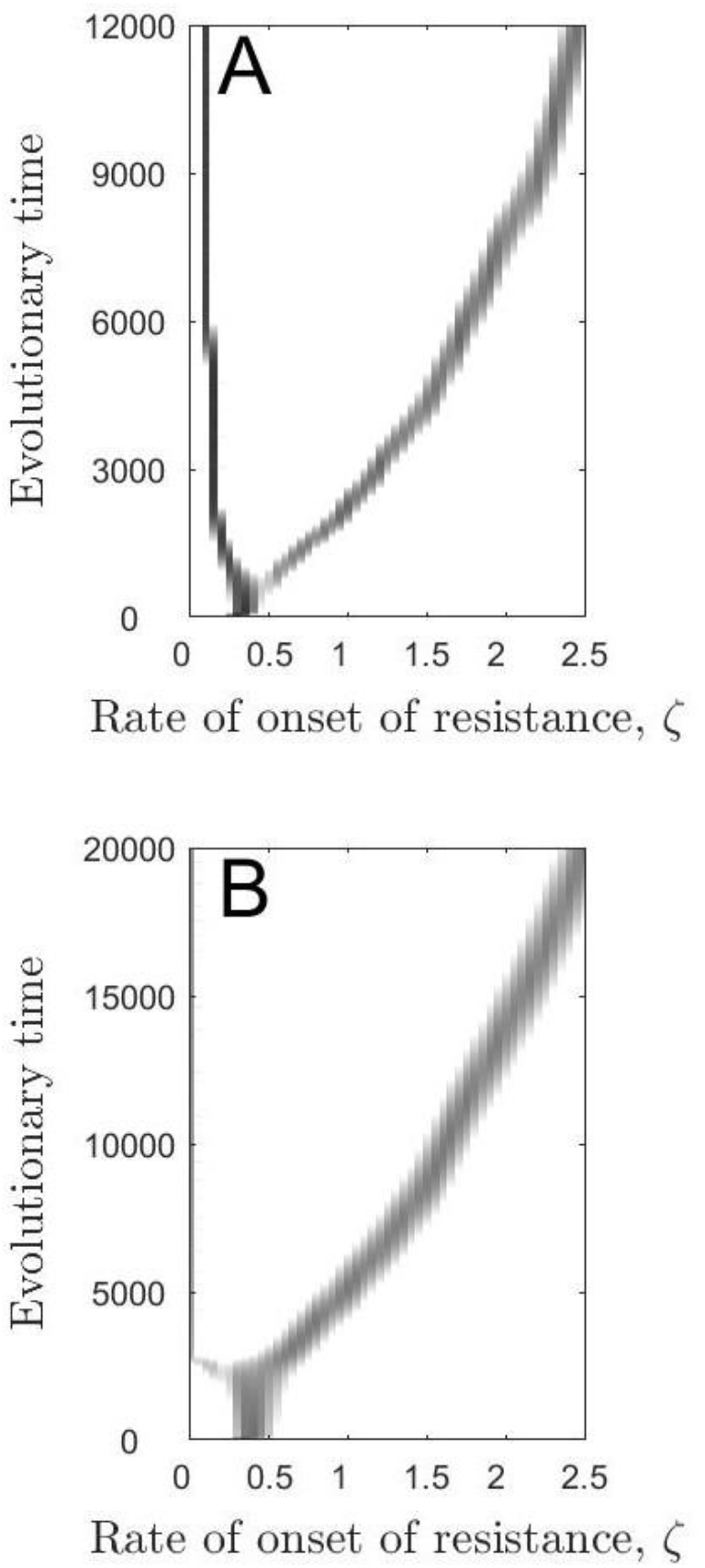
Simulated trajectories of the evolution of the rate of onset of resistance over time. (A) When costs of fast onset of resistance are paid throughout the lifetime of the host, the lower branch often tends towards a low, non-zero value whereas the upper branch goes to infinity. (B) When costs are paid only before the onset of resistance, the lower branch goes to zero whereas the upper branch goes to infinity. Parameters used are as in Table 1 with 1 – *f* = 0.5 and *α* = 0. We also use (a) 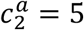 and (b) 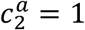 to generate branching points.

**Fig. 7.**
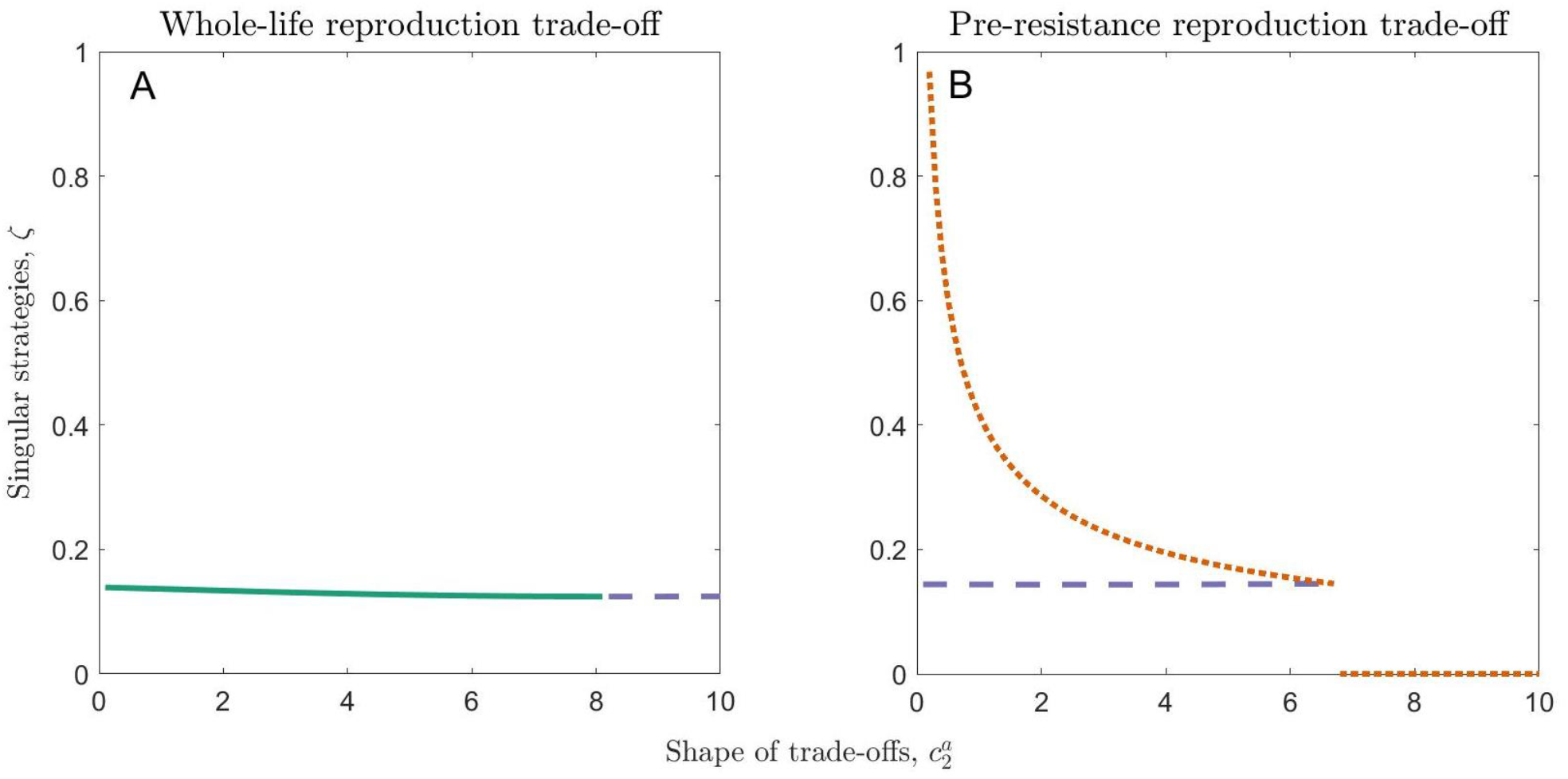
The effect of the shape of trade-offs on the incidence of branching points for different timings of costs. Continuously stable strategies are shown in green (solid), repellers are shown in orange (dotted) and branching points are shown in purple (dashed). (A) When costs of fast onset of resistance are paid throughout the host’s lifetime, costs must be strongly decelerating (high 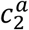) for branching to occur. (B) When costs are paid only before the onset of resistance, they must be weakly decelerating for branching to occur. Parameters used are as in Table 1 with 1 – *f* = 0 and *α* = 1.

When costs of having a faster onset of resistance are paid throughout the lifetime of the host, branching is most commonly observed when the baseline reproduction rate (*a*_0_), disease transmissibility (*β*) and sterility virulence (1 – *f*) are high, baseline mortality (*b*_0_) and the strength of trade-offs (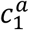 or 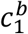) are low and mortality virulence (*a*) is intermediate (Fig. S1). When costs of fast onset of resistance are paid only before the onset of resistance, branching is most common when the baseline reproduction rate (*a*_0_), disease transmissibility (*β*), mortality virulence (*α*), baseline mortality rate (*b*_0_) and strength of the mortality trade-off 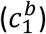 take intermediate values, when sterility virulence (1 – *f*) is high and when the strength of the reproduction trade-off 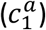 is low (Fig. S2).

## Discussion

Recent theory has shown how trade-offs and life-history traits combine to shape the evolution of age-specific resistance (Ashby and Bruns 2018; Buckingham et al. 2023). Here, we have shown how the average age at which hosts become resistant (1/*ζ*) evolves, depending on how and when costs of resistance are paid (costs only paid before or after the onset of resistance, or throughout the host’s lifetime). Our key findings are that (1) the rate of onset of resistance peaks for low mortality virulence, high sterility virulence and intermediate pathogen transmissibility; (2) when costs of resistance are paid throughout the lifetime of the host (or only after the onset of resistance) the onset of resistance typically evolves to a stable, intermediate rate whereas when costs are paid only before the onset of resistance, hosts are more likely to evolve no onset of resistance or an extremely fast onset of resistance (depending on parameters and initial conditions); and (3) pathogens may drive host diversification in the rate of onset of resistance.

Hosts evolve the fastest onset of resistance when pathogens cause high sterility virulence, low mortality virulence and intermediate transmissibility. Such pathogens induce the highest selection for resistance because they are able to maintain a significant density within the host population but also cause a large fitness cost to their hosts. Given a fixed, low-to-intermediate level of sterility virulence, an intermediate level of mortality virulence leads to the fastest rate of onset of resistance, but we have found that the combination of high sterility virulence and low mortality virulence leads to the fastest rate of onset overall in our model. This could be tested by determining the level of sterility or mortality inflicted by pathogens against which hosts experience a particularly early onset of resistance. We tentatively hypothesise that such pathogens would be more likely to inflict high levels of sterility virulence than high levels of mortality virulence.

The life-stage at which trade-offs occur has a significant effect on the evolution of the rate of onset of resistance. It is well established that the magnitude and shape of trade-offs, as well as the life-history traits involved, are critical to the evolution of various traits, including resistance (Kisdi 2006; Ashby and Bruns 2018; Buckingham et al. 2023). We have shown that the life-stage at which trade-offs act also drastically alters the evolutionary outcome for the rate of onset of resistance. An intermediate rate of onset of resistance is most likely to evolve when costs are paid throughout the host’s lifetime whereas very fast or very slow rates of onset, or dimorphism, are more likely if the costs are paid predominantly during the development of resistance. Very little empirical work has considered age-specific costs of resistance. However, we tentatively predict that organisms with very fast, very slow or polymorphic onset of resistance are most likely to experience costs which only act early in life.

When the rate of onset of resistance diversifies, it results in a fully (or almost fully) susceptible subpopulation coexisting with a fully resistant subpopulation. Such dimorphic populations, where some individuals are effectively resistant throughout their lives and others are always (or almost always) susceptible, are often seen in real, biological systems (Parker 1988; Jarosz and Burdon 1990; Wayne et al. 1996; Turkington et al. 2019). This evolutionary endpoint is very similar to classical models of host resistance evolution which result in dimorphic host populations with resistant and susceptible phenotypes (Antonovics and Thrall 1994; Boots and Haraguchi 1999). However, the mechanism for arriving at this endpoint is very different in our model. We have shown that this evolutionary endpoint can also arise through evolution of the rate of onset of resistance, where all hosts may transition from being susceptible to resistant as they age, rather than resistance/susceptibility being fixed before birth. This suggests that there may be an alternative evolutionary history of populations which are dimorphic in disease resistance, with the rate of onset of resistance diverging rather than the level of resistance. If this is indeed the case in some species, then one would expect “susceptible” hosts typically to have resistance genes that are suppressed rather than absent. However, it is possible that once the rate of onset of resistance approaches zero, resistance genes are lost rather than remaining suppressed. Experimental evolution may offer another path to testing this result by always selecting for hosts that have an earlier or later onset of resistance, thus validating whether a similar evolutionary endpoint can be achieved through the evolution of the timing of onset of resistance.

This paper constitutes an initial investigation into the theoretical consequences of the evolution of the age of onset of disease resistance. As such, we have considered a simple model which allows us to determine some possible outcomes of this evolutionary process. There are many additions and adjustments which could make this a more realistic model for specific biological systems. For example, we only focus on host evolution, but changes in patterns of host resistance are likely to affect pathogen evolution as well (Buckingham and Ashby 2022). We have also only considered the case of a disease with no recovery, as recovery would require differential handling of infection-acquired immunity or individuals who experience the onset of resistance while infected. Although this is a reasonable assumption for some host-pathogen interactions where recovery is negligible, such as systemic infections in plants, invertebrates, and bacteria (Ebert 2008; Abatángelo et al. 2017; Bruns et al. 2017), a more general development of the theory here would require careful consideration of how recovery affects immunity and post-infection sequelae. Recovery could be incorporated into our model in several different ways, depending on the mechanisms underlying recovery and the onset of resistance. For example, one could allow infected individuals to return to the susceptible class, but then the rate of onset of resistance takes on a different interpretation, as the process of becoming resistant is effectively turned off and on again as the host becomes infected and then later recovers. Alternatively, one could allow infected individuals to move into the resistant class if the onset of resistance occurred while they were infected, but would this process cause the host to clear the infection or just become resistant to re-infection? These scenarios cover rather different biological scenarios and so deserve careful consideration in future work.

For simplicity, we assumed that hosts experience a rapid onset of disease resistance (i.e. transitioning from susceptible to resistant in a short space of time). Rapid changes from low to high levels of resistance have been observed to occur in a number of biological systems. For example, Kurtis *et al*. showed that malaria (*Plasmodium falciparum*) resistance increases significantly in boys during puberty (between ages 15 and 20) which is relatively quick in relation to their lifespan (Kurtis et al. 2001). Critchlow *et al*. found that flour beetles (*Tribolium castaneum*) express more antimicrobial proteins as pupae than as larvae, resulting in a sudden increase in resistance during metamorphosis (Critchlow et al. 2019). Bruns *et al*. showed that alpine carnations (*Dianthus pavonius*) are far more resistant to anther-smut disease (*Microbotryum*) as adults than during the seedling stage, suggesting a rapid onset of resistance at maturation (Bruns et al. 2017). Farber & Mundt found that wheat plants (*Triticum aestivum*) significantly increase their resistance to stripe rust (*Puccinia striiformis*) between the ages of 3 and 5 weeks (with an average lifespan of 20 to 25 weeks) (Farber and Mundt 2017). As none of these changes in resistance occur instantaneously, however, our model could be extended to include not just completely susceptible and completely resistant stages, but also an additional stage in which resistance is intermediate, to represent the period during which these organisms are developing resistance. In cases where this stage is very short, our model should provide a good approximation.

However, some species do experience a very rapid onset of complete resistance, as in our model. For example, nucleopolyhedrovirus can only infect gypsy moths at the larval stage, with moths becoming fully resistant to the virus at the point of maturation (Elderd et al. 2008). This system closely mirrors our model and so may be a good candidate for empirical testing of the predictions made in this paper. Alternatively, anther-smut infections of Alpine carnations have previously been used as a model system for investigating age-specific differences in resistance (Bruns 2019). Seedlings are often significantly more susceptible to infection than adult plants and so this may provide an alternative system for testing our predictions empirically. To our knowledge, no empirical papers have sought to quantify the precise timing of the onset of resistance, or how this relates to characteristics of the disease.

Our model also assumes that hosts are born completely susceptible and later in life become fully resistant. Although large increases in the level of disease resistance may occur during a host’s lifetime, in many cases the host may shift between quantitative levels of resistance rather than qualitative resistance (Kurtis et al. 2001; Farber and Mundt 2017; Bruns et al. 2017; Critchlow et al. 2019). In future, our model should be extended to explore the timing of shifts between different levels of partial resistance. There are also empirical examples of reductions in resistance with host age and so the timing of the loss of resistance could be considered. For instance, Klemme *et al*. found that Atlantic salmon (*Salmo salar*) experience a reduction in resistance to a trematode (*Diplostomum pseudospathaceum*) when they mature from resident to migrant life-stages (Klemme et al. 2022). Similarly, species can also experience multiple changes in resistance during their lifetime. For instance, Garbutt *et al*. found that disease resistance in *Daphnia magna* increases as hosts mature into adults but subsequently decreases after their reproductive prime (Garbutt et al. 2014). A second change in resistance could be incorporated into our model by allowing resistant individuals to move into an older-age susceptible class, from which they could again become infected. This might reduce selection for a fast onset of resistance as the benefits of being resistant would be felt for a shorter time.

Overall, we have shown how the rate of onset of resistance evolves in response to variation in the nature of associated trade-offs and disease characteristics, potentially leading to host diversification and the same evolutionary endpoint as classical models of resistance evolution. Our model therefore provides an alternative explanation for the origin of dimorphism in host resistance.

## Data availability

The source code generated and analysed during the current study is available in the GitHub repository at: github.com/ecoevotheory/Buckingham_and_Ashby_2022b

