## Supplementary Materials for "The evolution of the age of onset of resistance to infectious disease"

Lydia J. Buckingham[1,2,*] and Ben Ashby[1,2,3,†]

1. Department of Mathematical Sciences, University of Bath, Bath, UK
2. Milner Centre for Evolution, University of Bath, Bath, UK
3. Department of Mathematics, Simon Fraser University, Burnaby, BC, Canada

### NON-DIMENSIONALISATION

We can scale the system of equations (1a) to (1c) in the main text as follows:

$$S, I, R, N \sim \frac{1}{q} \qquad (S1a)$$

$$t \sim \frac{1}{b_0} \qquad (S1b)$$

$$a, a_R, b, b_R \zeta \sim b_0 \qquad (S1c)$$

$$\beta \sim q b_0 \qquad (S1d)$$

which allows us to set $q = 1$ and $b_0 = 1$ without loss of generality.

### DERIVATION OF $R_0$

The dynamics of a small number of infected individuals in a population close to its disease-free equilibrium are given by:

$$\frac{\mathrm{d}I}{\mathrm{d}t} = \beta S^* I - b(1+\alpha)I \qquad (S2)$$

The disease can invade if $\frac{\mathrm{d}I}{\mathrm{d}t} > 0$ and so the basic reproductive ratio is given by:

$$R_0 = \frac{\beta S^*}{b(1+\alpha)} \qquad (S3)$$

### POPULATION VIABILITY

In the absence of disease, our system becomes:

$$\frac{dS}{dt} = a(1 - qN)S + a_R(1 - qN)R - bS - \zeta S \qquad (S4a)$$

$$\frac{dR}{dt} = -b_R R + \zeta S \qquad (S4b)$$

which has a single (non-trivial) stable equilibrium at:

$$S^* = \frac{b_R(ab_R + a_R\zeta - bb_R - b_R\zeta)}{q(b_R + \zeta)(ab_R + a_R\zeta)} \qquad (S5a)$$

$$R^* = \frac{\zeta}{b_R}S^* \qquad (S5b)$$

Hence, a disease-free population is viable if $ab_R + a_R\zeta > bb_R + b_R\zeta$.

### ENDEMIC EQUILIBRIUM

We can find the roots of the original system of equations (1a) to (1c). This tells us that:

$$S^* = \frac{b(1 + \alpha)}{\beta} \qquad (S6a)$$

$$R^* = \frac{\zeta}{b_R}S^* \qquad (S6b)$$

and

$$I^* = \frac{-B + \sqrt{B^2 - 4AC}}{2A} \qquad (S6c)$$

where

$$A = af \qquad (S6d)$$

$$B = afS^* + afR^* - af + \beta S^* + aS^* + a_R R^* \qquad (S6e)$$

$$C = bS^* + \zeta S^* - aS^* - a_R R^* + aS^{*2} + a_R S^* R^* + aS^* R^* + a_R R^{*2} \qquad (S6f)$$

Note that $R_0 > 1 \Leftrightarrow C < 0$ and so $I^* > 0$ whenever the basic reproductive ratio is greater than one.

We can linearise our model about this equilibrium to give the system:

$$\frac{\mathrm{d}}{\mathrm{d}t}\begin{pmatrix} S \\ I \\ R \end{pmatrix} = J\begin{pmatrix} S \\ I \\ R \end{pmatrix} \qquad (S7a)$$

where the Jacobian matrix is given by:

$$J = \begin{pmatrix} a(1-qN^*) - aq(S^*+fI^*) - a_R qR^* - \beta I^* - b - \zeta & af(1-qN^*) - aq(S^*+fI^*) - a_R qR^* - \beta S^* & a_R(1-qN^*) - aq(S^*+fI^*) - a_R qR^* \\ \beta I^* & \beta S^* - b(1+\alpha) & 0 \\ \zeta & 0 & -b_R \end{pmatrix} \quad (S7b)$$

The endemic equilibrium is linearly stable if the real part of each of the eigenvalues of $J$ is negative. We cannot calculate the eigenvalues of this matrix analytically and so we do this numerically for different sets of parameter values (see Matlab file "ecological_stability.m" in the source code). We find that the endemic equilibrium is unique and linearly stable whenever it exists, across a wide range of parameter values.

### INVASION FITNESS

The invasion dynamics of a rare host mutant with rate of onset of resistance $\zeta_m$ in an established resident population (at the endemic equilibrium, denoted by asterisks) are given by:

$$\frac{\mathrm{d}S_m}{\mathrm{d}t} = a(1-qN^*)(S_m + fI_m) + a_R(1-qN^*)R_m - \beta S_m I^* - bS_m - \zeta_m S_m \quad (S8a)$$

$$\frac{\mathrm{d}I_m}{\mathrm{d}t} = \beta S_m I^* - b(1+\alpha)I_m \quad (S8b)$$

$$\frac{\mathrm{d}R_m}{\mathrm{d}t} = -b_R R_m + \zeta_m S_m \quad (S8c)$$

Note that the mutant is assumed to be sufficiently rare as to make the effect of mutant-mutant interactions negligible and so these interactions are not included in the system above.

In the cases where $a$, $a_R$, $b$ or $b_R$ trade off with the rate of resistance onset, $\zeta$, these terms are functions of $\zeta_m$ in the equations above.

The invasion fitness is derived using the next-generation method (Hurford et al., 2010), with the following decomposition of the Jacobian matrix, $A$:

$$A = \begin{pmatrix} a(1-qN^*) & a(1-qN^*)f & a_R(1-qN^*) \\ 0 & 0 & 0 \\ 0 & 0 & 0 \end{pmatrix} - \begin{pmatrix} \beta I^* + b + \zeta_m & 0 & 0 \\ -\beta I^* & b(1+\alpha) & 0 \\ -\zeta_m & 0 & b_R \end{pmatrix} \quad (S9)$$

Denoting this decomposition as $A = F - V$, the invasion fitness is one less than the spectral radius of $FV^{-1}$:

$$w(\zeta_m, \zeta) = \frac{a(1-qN^*)b(1+\alpha)b_R + a(1-qN^*)f\beta I^* b_R + a_R(1-qN^*)b(1+\alpha)\zeta_m}{(\beta I^* + b + \zeta_m)b(1+\alpha)b_R} - 1 \qquad (S10)$$

Note that $I^*$ and $N^*$ will typically be functions of $\zeta$ and, depending on which trade-offs are being considered, $a$, $a_R$, $b$ or $b_R$ may be functions of $\zeta_m$.

### Ecological system for arbitrary number of phenotypes

In our evolutionary simulations, we relax the assumption of rare mutations so that multiple mutant phenotypes may exist alongside the resident at any given time. Also, the resident may not be at its ecological equilibrium by the time new mutants appear. To do this, we need to consider the ecological system for an arbitrary number of host strains. Let $S_i$, $I_i$ and $R_i$ denote the density of susceptible, infected and resistant individuals of strain $i$ for $1 \leq i \leq n$ and let $\zeta_i$ be the rate of onset of resistance for strain $i$. Then the ecological dynamics are given by:

$$\frac{dS_i}{dt} = a\big(1-q\hat{N}\big)(S_i + fI_i) + a_R\big(1-q\hat{N}\big)R_i - \beta S_i\hat{I} - bS_i - \zeta_i S_i \qquad (S11a)$$

$$\frac{dI_i}{dt} = \beta S_i \hat{I} - b(1+\alpha)I_i \qquad (S11b)$$

$$\frac{dR_i}{dt} = -b_R R_i + \zeta_i S_i \qquad (S11c)$$

where $\hat{N} := \sum_{i=1}^{n}(S_i + I_i + R_i)$ is the total population density and $\hat{I} := \sum_{i=1}^{n} I_i$ is the total infected density. Note that when $a$, $a_R$, $b$ or $b_R$ are functions of the evolving trait, they are functions of $\zeta_i$ in the strain $i$ equations.

### Constant costs

#### *Singular strategies*

Singular strategies are turning points of the invasion fitness function, calculated by differentiating the invasion fitness ($w$) with respect to the mutant trait, $\zeta_m$, and then finding the roots of this fitness gradient in the case where the resident and mutant traits are equal ($\zeta_m = \zeta$).

In scenarios (3) to (6), where $a$, $a_R$, $b$ or $b_R$ are functions of $\zeta$, it is not possible to write down a closed-form expression for the singular strategy. However, this is possible in scenarios (1) and (2), where $a_R = ah$, $b_R = b(1+\delta)$ and $a$ and $b$ are constants (note that $\delta = 0$ in scenario (1) and $h = 1$ in scenario (2)).

In this case, the fitness gradient may be written as:

$$\frac{\partial w}{\partial \zeta_m}\Big|_{\zeta_m=\zeta} = \frac{a_R(1-qN^*) - b_R}{(\beta I^* + b + \zeta)b_R} \qquad (S12)$$

where $I^*$ and $N^*$ are functions of $\zeta$.

We can combine this equation with the original system of equations (1a) to (1c) in the main text. Finding the roots of these four equations simultaneously allows us to calculate the singular strategy ($\zeta^*$) and the endemic equilibrium at the singular strategy, as follows:

$$\zeta^* = b(1+\delta)\left(\frac{p\ -\ 1}{fp\ -\ (1+\alpha)} + \frac{\beta(a\ -\ bp)}{aq\ b(1+\alpha)} -\ 1\right) \qquad (S13a)$$

$$S^*|_{\zeta=\zeta^*} = \frac{b(1+\alpha)}{\beta} \qquad (S13b)$$

$$R^*|_{\zeta=\zeta^*} = \frac{\zeta^*}{b_R}S^* \qquad (S13c)$$

$$I^*|_{\zeta=\zeta^*} = \frac{p-1}{(1+\alpha)-fp}S^* \qquad (S13d)$$

where $p = \frac{1+\delta}{h}$ for notational convenience.

In the absence of disease (and the presence of costs of resistance), the rate of onset of resistance will always evolve to become zero (no onset of resistance). We can see that $I^*|_{\zeta=\zeta^*} > 0 \Leftrightarrow (1+\alpha) > fp$ and so the onset of resistance can only evolve if $h(1+\alpha) > f(1+\delta)$. This condition represents the costs of infection being sufficiently higher than the costs of resistance.

*INVASION FITNESS*

We would like to know the invasion fitness of a rare mutant within a resident population at the singular strategy, $w(\zeta_m, \zeta^*)$.

We saw above from equation S12 that, at the singular strategy, $a_R(1-qN^*) - b_R = 0$ must be satisfied. Substituting this into the expression for the invasion fitness (equation S10) tells us that:

$$w(\zeta_m, \zeta^*) = \frac{\beta I^* - \dfrac{b(1+\alpha)(p-1)}{(1+\alpha)-fp}}{\left(\dfrac{(\beta I^* + b + \zeta_m)^2(1+\alpha)}{(1+\alpha)-fp}\right)} \qquad (S14)$$

where $I^*$ is evaluated at the singular strategy. We know from equation S13d that $I^*|_{\zeta=\zeta^*} = \frac{(p-1)b(1+\alpha)}{((1+\alpha)-fp)\beta}$ and so we can see that $w(\zeta_m, \zeta^*) \equiv 0$.

Therefore, any derivative of $w(\zeta_m\ , \zeta)$ taken only with respect to $\zeta_m$ and subsequently evaluated at $\zeta = \zeta^*$ will be equal to zero (including the condition for evolutionary stability).

*CONVERGENCE STABILITY*

Since we already know that $\frac{\partial^2 w}{\partial \zeta_m^2}|_{\zeta_m=\zeta=\zeta^*} = 0$, a singular strategy will be convergence stable if and only if $\frac{\partial^2 w}{\partial \zeta_m \partial \zeta}|_{\zeta_m=\zeta=\zeta^*} < 0$. Due to the complexity of the general expression for the endemic equilibrium (equation S6c), we cannot write down an expression for $\frac{\partial^2 w}{\partial \zeta_m \partial \zeta}|_{\zeta_m=\zeta=\zeta^*}$ which is simple enough for us to determine its sign.

The Matlab file "constant_costs_convergence_stability.m" (included in the source code) determines the sign of this term numerically for many sets of parameter values. It reveals that the singular strategy is convergence stable if and only if $h(1+\alpha) > f(1+\delta)$.

We have already shown that there cannot be a positive singular strategy unless this condition holds. Therefore, whenever there is a singular value of the rate of onset of resistance, this singular strategy is convergence stable.

**FAST ONSET OF RESISTANCE**

We have found that the endemic equilibrium of our original system satisfies:

$$S^* = \frac{b(1+\alpha)}{\beta} \qquad (S15a)$$

$$R^* = \frac{\zeta}{b_R} S^* \qquad (S15b)$$

and

$$A{I^*}^2 + BI^* + C = 0 \qquad (S15c)$$

where

$$A = af \qquad (S15d)$$

$$B = afS^* + afR^* - af + \beta S^* + aS^* + a_R R^* \qquad (S15e)$$

$$C = bS^* + \zeta S^* - aS^* - a_R R^* + a{S^*}^2 + a_R S^* R^* + aS^* R^* + a_R {R^*}^2 \qquad (S15f)$$

As $\zeta \to \infty$, $C \sim \frac{a_R {S^*}^2}{b_R^2}\zeta^2 + O(\zeta)$ and $B \sim \frac{(af + a_R)S^*}{b_R}\zeta + O(1)$ and $A \sim O(1)$.

This means that equation S15c can only balance if $I^* \to -\infty$ as $\zeta \to \infty$. Clearly, this is not possible (the number of infected individuals must always be non-negative). Therefore, there is no endemic equilibrium when $\zeta$ is sufficiently large. That is, $I^* = 0$ when $\zeta$ is large.

Taking $I^* = 0$ allows us to determine the ecological equilibrium for scenarios (3) and (4):

$$S^* = \frac{b(a-b)}{a(b+\zeta)} \qquad (S16a)$$

$$R^* = \frac{\zeta(a-b)}{a(b+\zeta)} \qquad (S16b)$$

and for scenarios (5) and (6):

$$S^* = \frac{b_0(ab_0 + a_0\zeta - bb_0 - b_0\zeta)}{(b_0 + \zeta)(ab_0 + a_0\zeta)} \qquad (S17a)$$

$$R^* = \frac{\zeta(ab_0 + a_0\zeta - bb_0 - b_0\zeta)}{(b_0 + \zeta)(ab_0 + a_0\zeta)} \qquad (S17b)$$

We can also simplify the invasion fitness (equation S10) in each of these scenarios and, using the above expressions for the ecological equilibrium, derive the fitness gradient in scenarios (3) and (4):

$$\frac{\partial w}{\partial \zeta_m}|_{\zeta_m=\zeta} = \frac{1}{a}\frac{\mathrm{d}a}{\mathrm{d}\zeta} - \frac{1}{b}\frac{\mathrm{d}b}{\mathrm{d}\zeta} \qquad (S18)$$

and in scenarios (5) and (6):

$$\frac{\partial w}{\partial \zeta_m}|_{\zeta_m=\zeta} = \frac{a_0 b - ab_0}{(b + \zeta)(ab_0 + a_0\zeta)} + \frac{b_0}{ab_0 + a_0\zeta}\frac{\mathrm{d}a}{\mathrm{d}\zeta} - \frac{1}{b + \zeta}\frac{\mathrm{d}b}{\mathrm{d}\zeta} \qquad (S19)$$

We know that $\frac{\mathrm{d}a}{\mathrm{d}\zeta} < 0$ and $\frac{\mathrm{d}b}{\mathrm{d}\zeta} > 0$ and so we can see that $\frac{\partial w}{\partial \zeta_m}|_{\zeta_m=\zeta} < 0$ for sufficiently large $\zeta$ in scenarios (3) and (4). This means that the uppermost singular strategy will always be an evolutionary attractor in these scenarios (it is convergence stable).

We also know that $a_0 > a$ and $b_0 < b$ and that $\frac{\mathrm{d}a}{\mathrm{d}\zeta}$ and $\frac{\mathrm{d}b}{\mathrm{d}\zeta}$ decay exponentially to zero as $\zeta$ increases when $c_2^i > 0$. However, when $c_2^i < 0$, we know that $\frac{\mathrm{d}a}{\mathrm{d}\zeta} \to -\infty$ and $\frac{\mathrm{d}b}{\mathrm{d}\zeta} \to \infty$ as $\zeta$ increases. Therefore, $\frac{\partial w}{\partial \zeta_m}|_{\zeta_m=\zeta} > 0$ for sufficiently large $\zeta$ in scenarios (5) and (6) when $c_2^a$ and $c_2^b$ are positive whereas $\frac{\partial w}{\partial \zeta_m}|_{\zeta_m=\zeta} < 0$ for sufficiently large $\zeta$ in scenarios (5) and (6) when $c_2^a$ and $c_2^b$ are negative.

This means that the uppermost singular strategy will always be a repeller (convergence unstable) when $c_2^a$ and $c_2^b$ are positive and will always be an attractor when $c_2^a$ and $c_2^b$ are negative, in scenarios (5) and (6).

### DIMORPHIC POPULATIONS

We consider a dimorphic population where one sub-population has no onset of resistance ($\zeta = 0$). This system is represented by the equations:

$$\frac{\mathrm{d}S_1}{\mathrm{d}t} = a_0(1 - qN)(S_1 + fI_1) - \beta S_1 I - b_0 S_1 \qquad (S20a)$$

$$\frac{\mathrm{d}I_1}{\mathrm{d}t} = \beta S_1 I - b_0(1+\alpha)I_1 \qquad (S20b)$$

$$\frac{\mathrm{d}S_2}{\mathrm{d}t} = a(1-qN)(S_2 + fI_2) + a_R(1-qN)R_2 - \beta S_2 I - bS_2 - \zeta S_2 \qquad (S20c)$$

$$\frac{\mathrm{d}I_2}{\mathrm{d}t} = \beta S_2 I - b(1+\alpha)I_2 \qquad (S20d)$$

$$\frac{\mathrm{d}R_2}{\mathrm{d}t} = -b_R R_2 + \zeta S_2 \qquad (S20e)$$

where $I := I_1 + I_2$ and $N := S_1 + I_1 + S_2 + I_2 + R_2$.

We seek to determine the evolutionary dynamics of the rate of onset of resistance of the second sub-population. As we have only observed dimorphism in the case of trade-offs with $c_2^i > 0$, we only consider the system in this case.

We have observed from simulations that the rate of onset of resistance, $\zeta$, of the second sub-population increases significantly over evolutionary time. We wish to determine whether $\zeta$ will evolve to increase indefinitely. To do this, we consider the dynamics of the above system as $\zeta \to \infty$.

By setting each of the above equations equal to zero, we can determine the ecological equilibrium of the system. In scenarios (5) and (6), where $a_R = a_0$, $b_R = b_0$ and one of $a$ or $b$ varies with $\zeta$, we let $q = 1$ and $b_0 = 1$ (by non-dimensionalising) and find that

$$N^* \sim 1 - \frac{1}{a_0} - \frac{c_1^a}{a_0\left(1 - e^{-c_2^a}\right)\zeta} + O\left(\frac{1}{\zeta^2}\right) \qquad \text{as } \zeta \to \infty \qquad (S21a)$$

in the case of scenario (5) and

$$N^* \sim 1 - \frac{1}{a_0} - \frac{c_1^b}{a_0\left(1 - e^{-c_2^b}\right)\zeta} + O\left(\frac{1}{\zeta^2}\right) \qquad \text{as } \zeta \to \infty \qquad (S21b)$$

in the case of scenario (6).

The fitness gradient in either case is given by:

$$\frac{\partial w}{\partial \zeta_m}\Big|_{\zeta_m=\zeta} = \frac{a_0(1-N^*) - 1}{(\beta I^* + b + \zeta)} + \frac{\mathrm{d}a}{\mathrm{d}\zeta}\frac{(\beta I^* + b + \zeta) - a_0(1-N^*)\zeta}{(\beta I^* + b + \zeta)a} + \frac{\mathrm{d}b}{\mathrm{d}\zeta}\frac{a(1-N^*) + a_0(1-N^*)\zeta - b - (\beta I^* + b + \zeta)}{(\beta I^* + b + \zeta)b} \qquad (S22)$$

This expression satisfies:

$$\frac{\partial w}{\partial \zeta_m}\Big|_{\zeta_m=\zeta} \sim \frac{c_1^i}{\left(1 - e^{-c_2^i}\right)\zeta^2} + O\left(\frac{1}{\zeta^3}\right) \qquad \text{as } \zeta \to \infty \qquad (S23)$$

where $i = a$ in scenario (5) and $i = b$ in scenario (6).

This means that the fitness gradient is positive for sufficiently large values of $\zeta$ when $c_2^i > 0$. Therefore, when $c_2^i > 0$, we would expect $\zeta$ to evolve to increase indefinitely in the cases where simulations show that it rises significantly.

### DESCRIPTION OF EVOLUTIONARY SIMULATIONS

1. Run the ecological dynamics of the system for a fixed length of time, with the rate of onset of resistance taking its resident value.
2. Introduce a mutant, randomly determining whether the trait will mutate to be slightly higher or lower than its current value. Add a small sub-population with the new, mutant trait value.
3. Run the ecological dynamics of the system for a fixed length of time, starting at its current composition.
4. Remove any sub-populations which have a density below a certain threshold (they are extinct).
5. Introduce a mutant by randomly determining which sub-population (rate of onset of resistance trait value) the mutant will come from and whether the trait will mutate to be slightly higher or lower than its current value. Add a small sub-population with the new, mutant trait value.
6. Repeat steps 3 to 5 for many evolutionary timesteps.

In these simulations, ecological and evolutionary timescales are not completely separated because the ecological system does not necessarily reach its equilibrium before the next mutant is introduced. Mutations do not have arbitrarily small phenotypic effects because mutations alter the rate of onset of resistance by a small, fixed interval.

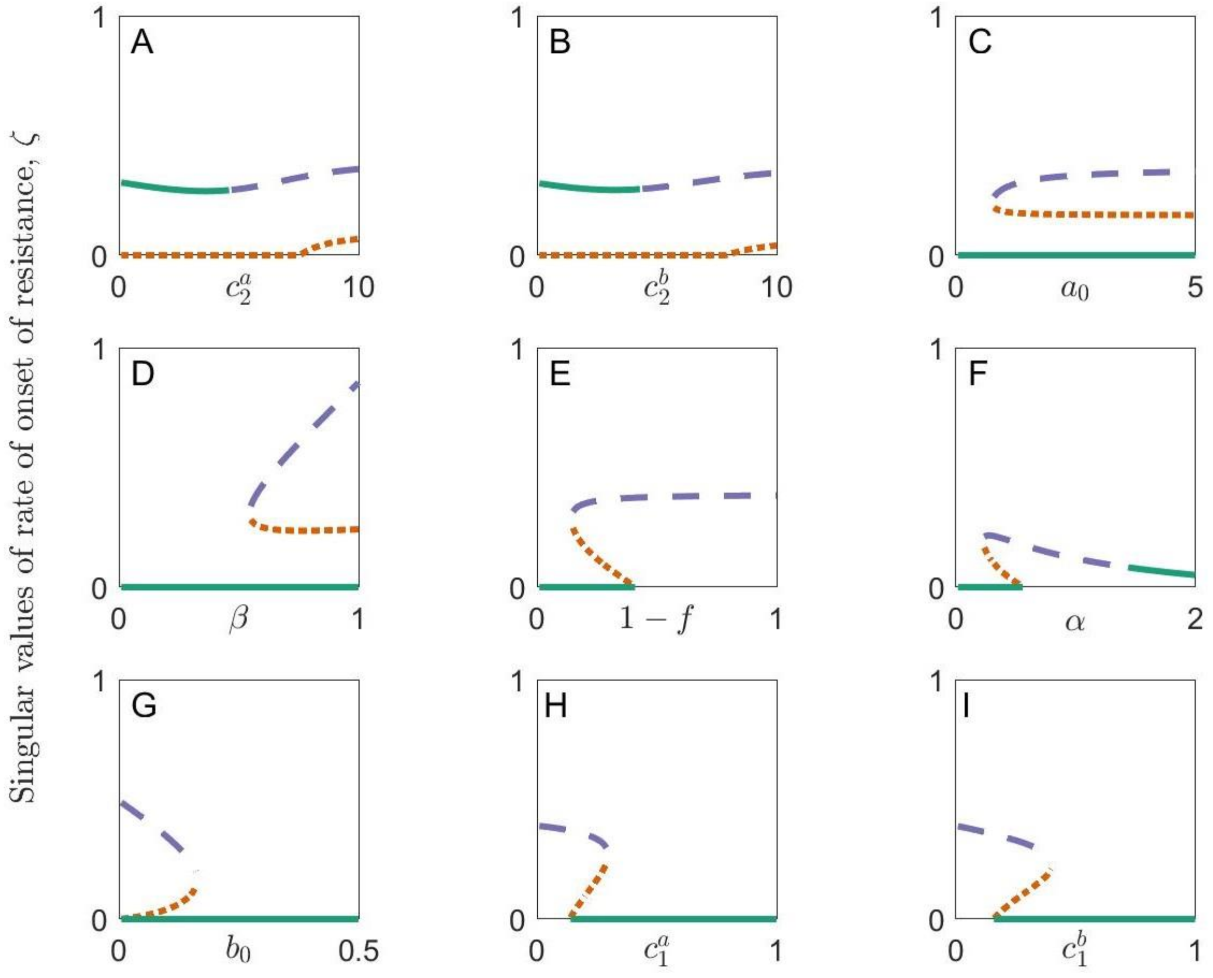


**Fig. S1** The effect of different parameters on the number and stability of singular strategies in the cases where costs of fast onset of resistance are paid throughout the lifetime of the host (case 3 for (A), (C), (E) & (H) and case 4 for (B), (D), (F), (G) & (I)). Continuously stable strategies are shown in green (solid), repellers are shown in orange (dotted) and branching points are shown in purple (dashed). Parameters used are as in Table 1 except for (A) $c_1^a = 0.18, 1-f = 0.5, \alpha = 0$, (B) $c_1^b = 0.2, 1-f = 0.5, \alpha = 0$, (C) $c_2^a = 10, 1-f = 0.2, \alpha = 0$, (D) $c_1^b = 0.2, c_2^b = 10, 1-f = 0.2, \alpha = 0$, (E) $c_2^a = 10, \alpha = 0$, (F) $c_2^b = 10, 1-f = 0$, (G) $c_1^b = 0.2, c_2^b = 10, 1-f = 0.5, \alpha = 0$, (H) $c_2^a = 10, 1-f = 0.5, \alpha = 0$ and (I) $c_2^b = 10, 1-f = 0.5, \alpha = 0$. Changing these parameters causes quantitative shifts in these figures but the qualitative patterns are consistent for a wide range of parameters.

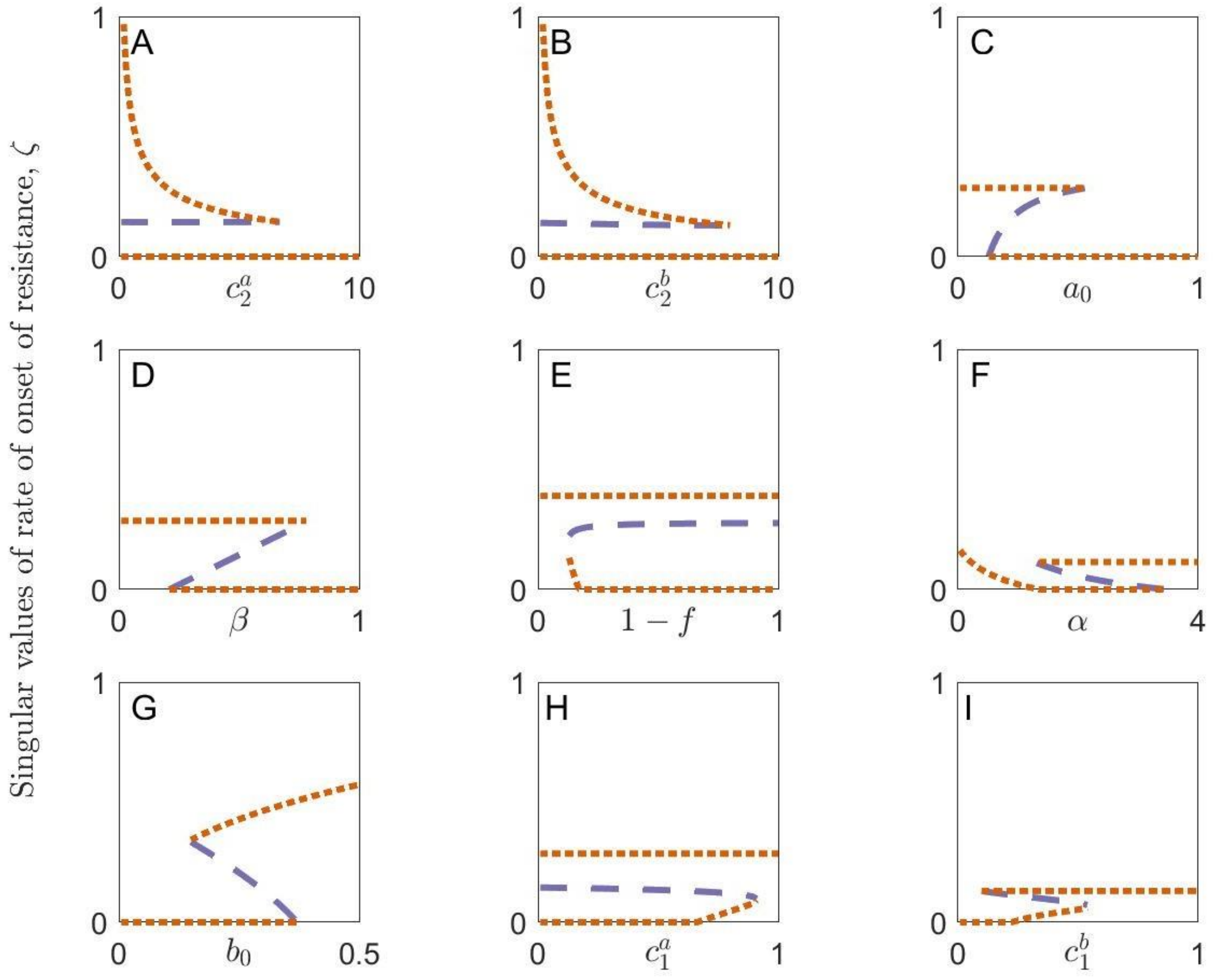


**Fig. S2** The effect of different parameters on the number and stability of singular strategies in the cases where costs of fast onset of resistance are paid only before the onset of resistance (case 5 for (A), (D), (E), (F) & (G) and case 6 for (B), (C) & (H)). Continuously stable strategies are shown in green (solid), repellers are shown in orange (dotted) and branching points are shown in purple (dashed). Parameters used are as in Table 1 except for (A) $1 - f = 0, \alpha = 1$, (B) $1 - f = 0, \alpha = 1$, (C) $1 - f = 0.5, \alpha = 0$, (D) $1 - f = 0, \alpha = 1$, (E) $b_0 = 0.2, \alpha = 0$, (F) $c_1^a = 0.2, c_2^a = 10, 1 - f = 0$, (G) $1 - f = 0.5, \alpha = 0$, (H) $1 - f = 0, \alpha = 1$ and (I) $c_2^b = 8, 1 - f = 0, \alpha = 1$. Changing these parameters causes quantitative shifts in these figures but the qualitative patterns are consistent for a wide range of parameters.
